# *PpRPK2* modulates auxin homeostasis and transport to specify stem cell identity and plant shape in the moss Physcomitrella

**DOI:** 10.1101/2021.06.24.449551

**Authors:** Zoe Nemec Venza, Connor Madden, Amy Stewart, Wei Liu, Ondřej Novák, Aleš Pěnčík, Andrew C. Cuming, Yasuko Kamisugi, C. Jill Harrison

**Affiliations:** School of Biological Sciences, University of Bristol, 24 Tyndall Avenue, Bristol, BS8 1TQ, UK; Division of Psychological Medicine & Clinical Neurosciences, MRC Centre for Neuropsychiatric Genetics & Genomics, Cardiff University School of Medicine, Heath Park, Cardiff, CF14 4XN, UK; Laboratory of Growth Regulators, Faculty of Science of Palacký University and Institute of Experimental Botany of the Czech Academy of Sciences, Šlechtitelů 27, 78371 Olomouc, Czech Republic; Centre for Plant Sciences, Faculty of Biological Sciences, University of Leeds, Leeds, LS2 9JT, UK

## Abstract

Plant shape is determined by the activity of stem cells in the growing tips, and evolutionary changes in shape are linked to changes in stem cell function. The CLAVATA pathway is a key regulator of stem cell function in the multicellular shoot tips of Arabidopsis, acting via the WUSCHEL transcription factor to modulate hormone homeostasis. Broad scale evolutionary comparisons have shown that CLAVATA is a conserved regulator of land plant stem cell function, but CLAVATA acts independently of WUSCHEL-like (WOX) proteins in bryophytes, raising questions about the evolution of stem cell function and the role of the CLAVATA pathway.

Here we show that the moss (Physcomitrella) CLAVATA pathway affects stem cell activity and overall plant shape by modulating hormone homeostasis. CLAVATA pathway components are expressed in the tip cells of filamentous tissues, regulating cell identity, filament branching patterns and plant spread. The PpRPK2 receptor-like kinase plays the major role and is expressed more strongly than other receptor-encoding genes. *Pprpk2* mutants have abnormal responses to cytokinin, and auxin transport inhibition and show reduced *PIN* auxin transporter expression.

We propose a model whereby *PpRPK2* modulates *PIN* activity to determine stem cell identity and overall plant form in Physcomitrella. Our data indicate that CLAVATA-mediated auxin homeostasis is a fundamental property of plant stem cell function likely exhibited by the last shared common ancestor of land plants.

## Introduction

Organ size and shoot architecture are determined by the number and activity of stem cells in the growing shoot tips of flowering plants such as Arabidopsis [1]. The size and integrity of Arabidopsis shoot tips is maintained by the action of a molecular feedback loop involving CLAVATA peptides and receptors and the WUSCHEL transcription factor [2, 3]. *CLAVATA3* is expressed in stem cells in the outermost cell layers [4], encoding a protein that is processed to form a small diffusible peptide [5–8]. The CLAVATA3 peptide acts as a ligand to the CLAVATA1 receptor which is active in inner cell layers of the shoot tip [9, 10], and signaling via CLAVATA1 confines the expression of *WUSCHEL* to a few cells at the centre of the shoot tip [2]. In turn, the WUSCHEL protein moves to the outermost cell layers of the shoot tip [11, 12], promoting expression of *CLAVATA3* [13], and mutants with defective CLAVATA or WUSCHEL function respectively over-proliferate cells in the shoot tips or are unable to maintain the stem cell population resulting in enlargement of the tips or shoot termination [9, 14]. The regulation of meristem function by CLAVATA depends on the maintenance of low levels of auxin signaling permissive to stem cell activity in the central zone of the shoot tips by WUSCHEL [2, 15].

Unlike Arabidopsis, the growing tips of mosses comprise a single apical cell [16]. Spores germinate to form a branching mat of filamentous tissue termed the protonema. When they first start to grow, protonemal tissues serve a mainly photosynthetic function, and the filaments have so called chloronemal identity, comprising short cells with dark green chloroplasts [17]. Tip growth in the apical cells of each filament extends plant spread, and the apical cells cleave in a plane perpendicular to the main axis of growth [18]. Later on, more rapidly growing filaments termed caulonemata develop, and caulonemal apical cells cleave obliquely to generate long cells with more numerous but less pigmented chloroplasts than in chloronemata [17]. The relative growth of chloronemata and caulonemata determines the shape of the plant, such that plants comprising solely chloronemata are small and round, whereas plants with predominantly caulonemata are larger and have an irregular foraging fringe [19].

Auxin, auxin transport, auxin perception and auxin signaling via a TIR1/AFB and AUX/IAA-dependent mechanism promote and are required for caulonemal differentiation [19–22]. In contrast, auxin resistant *ppafb* RNAi and *ppiaa2-P328S* mutants, miR1219 and tasiARF degradation-resistant *PpARFb2* and *PpARFb4* mutants, and plants treated with an auxin synthesis inhibitor (L-Kynurenine) cannot undergo the chloronema to caulonema developmental transition [19, 21, 23]. Protonemal apical cells are the site of cell fate decisions that affect overall plant shape, yet auxin reporters suggest that they have minimal auxin sensing [22]. Exogenously applied cytokinin has a converse effect to auxin, suppressing caulonemal differentiation. Cytokinin promotes AUX/IAA expression, and cytokinin-induced expression depends on auxin signaling [17, 19]. Thus, a complex interplay between auxin and cytokinin regulates protonemal tip cell identity, the chloronema to caulonema transition and overall plant shape.

The *Physcomitrella* (*Physcomitrium patens*) CLAVATA pathway comprises 9 CLAVATA3-like peptides (PpCLEs 1-9), 2 CLAVATA1-like receptors (PpCLV1a and PpCLV1b), and a further receptor similar to Arabidopsis RPK2 (PpRPK2) [1, 24, 25]. By RT-PCR and promoter fusions, previous work showed that *PpCLEs 1*, *2* and *7* and *PpCLV1a*, *PpCLV1b* and *PpRPK2* are expressed highly during the development of the multicellular haploid phase of the life cycle (in the gametophyte) and are required to establish stem cells that iterate shoot-like structures (gametophores), affecting stem cell growth, identity and division plane orientations [25]. There are 3 *P. patens WUSCHEL*-like homeobox (*WOX*) genes, *PpWOX13LA*, *PpWOX13LB* and *PpWOX13LC* [26]. By RT-PCR previous work detected constitutive expression of *PpWOX13LA*, *PpWOX13LB*, but no expression of *PpWOX13LC* [26] and protein fusions for PpWOX13LA and PpWOX13LB showed elevated expression in protonemal stem cells and stem cells forming in a leaf regeneration assay [26]. *Ppwox13lab* mutants were unable to initiate growth in leaf regeneration assays but were otherwise indistinguishable from wild-type plants during gametophyte development [26]. *Marchantia* CLAVATA pathway components (*MpCLV3* and *MpCLV1*) similarly act in a WOX-independent manner [27]. Thus, whilst CLAVATA is a conserved regulator of land plant stem cell function, WOX function is inessential in bryophytes, raising questions about the regulation of bryophyte stem cell function, the evolution of the *CLAVATA*-*WUSCHEL* gene regulatory network and regulators of stem cell function in the last shared common ancestor of land plants. Here we show that CLAVATA represses the chloronema to caulonema developmental transition and regulates plant spread and shape by modulating auxin homeostasis and PIN-mediated auxin transport.

## Results

### CLAVATA pathway components are expressed in protonemata and *PpRPK2* is the most strongly expressed receptor-encoding gene

To investigate roles for CLAVATA in protonemal development, we first engineered *promoter∷NLSGFPGUS* (*promoter∷NGG*) reporter lines for *PpCLEs* whose expression was uncharacterized in previous work, namely *PpCLE3*, *PpCLE4*, *PpCLE5*, *PpCLE6*, *PpCLE8* and *PpCLE9* (Figure 1A, Figure S1). Analysis of three- to four- week old plants revealed signal in protonemal tissues and gametophores for all *promoter∷NGG* lines, and expression patterns were consistent between independently generated lines (not shown). To determine when *CLAVATA* expression first arises during development, spores from previously established *PpCLE1∷NGG*, *PpCLE2∷NGG*, *PpCLE7∷NGG*, *PpCLV1a∷NGG*, *PpCLV1b∷NGG* and *PpRPK2∷NGG* lines [25] and newly engineered *PpCLE3∷NGG*, *PpCLE4∷NGG*, *PpCLE5∷NGG*, *PpCLE6∷NGG*, *PpCLE8∷NGG* and *PpCLE9∷NGG* lines were germinated and grown for 2 to 11 days prior to staining (Figure 1B, C). Whilst no signal was detected in any line at germination (Figure S2), *PpCLE3∷NGG*, *PpCLE9∷NGG* and *PpCLV1a∷NGG* plants comprising primary chloronemata showed expression (Figure 1B). Following the differentiation of caulonemata, expression was activated in the tip cells of all lines and was more intense and more frequently observed in caulonemal rather than chloronemal tip cells (Figure 1C).

**Figure 1:**
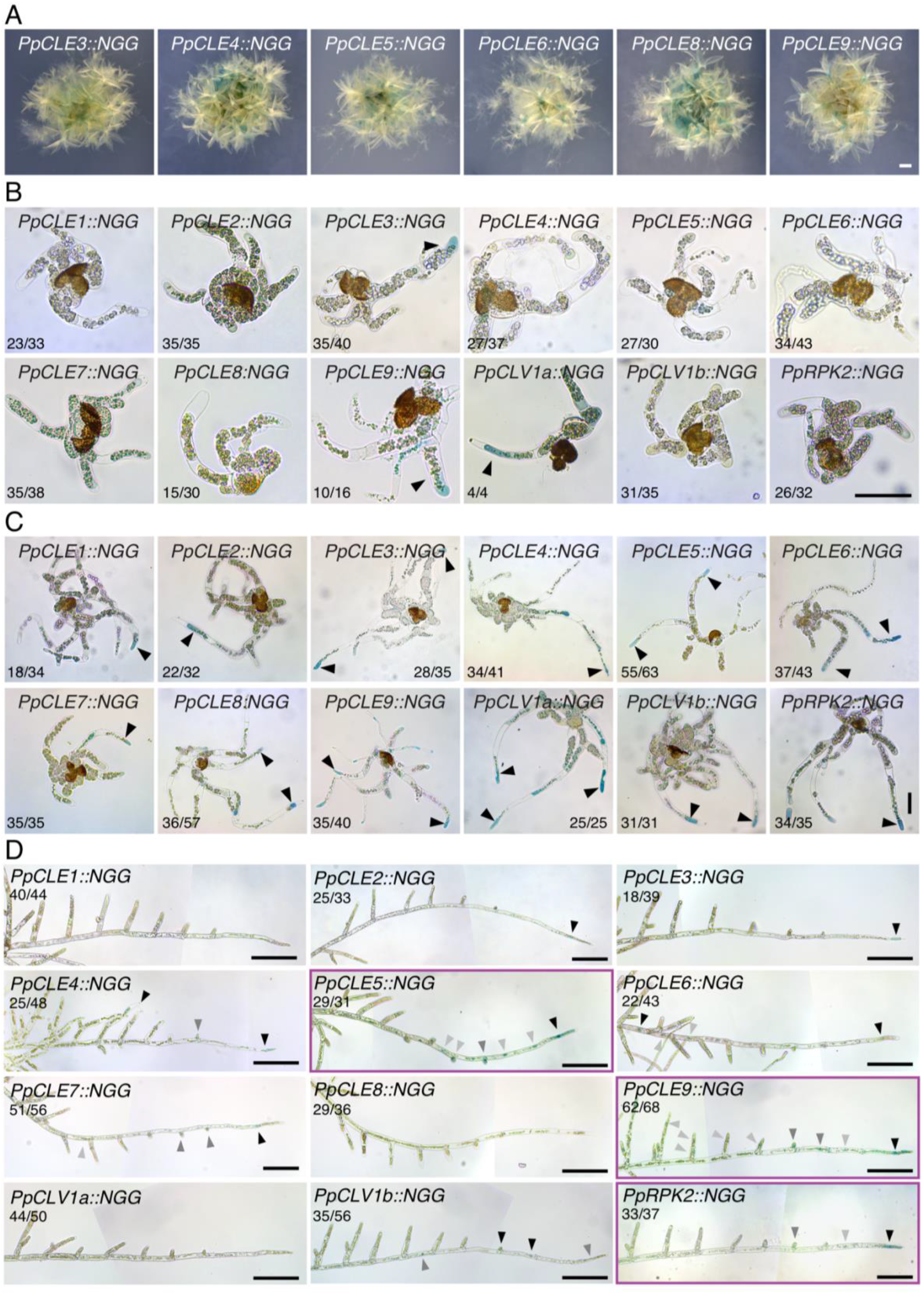
Caulonemal tip cells are likely sites of CLE peptide production and perception. **(A)** Micrographs showing expression patterns of newly generated *PpCLE∷NGG* lines in whole plants, showing signal in gametophores and protonemal tissues. Scale bar = 1 mm. **(B)** Micrographs of sporelings which comprised mainly chloronemata. *PpCLE3∷NGG*, *PpCLE9∷NGG* and *PpCLV1a∷NGG* lines showed expression. The numbers in each panel indicate the proportion of sporelings displaying a similar expression pattern. Scale bar = 50 μm. **(C)** Micrographs of sporelings comprising a mix of chloronemata and caulonemata. All lines showed *CLAVATA* expression. The signal was absent or weak in chloronemal cells, but stronger and more frequently detected in caulonemal tip cells (Black arrowheads). The numbers in each panel indicate the proportion of sporelings displaying a similar expression pattern. Scale bar = 50 μm. **(D)** Stitched light micrographs showing promoter∷NGG reporter expression in foraging filaments. Although no signal was detected in the majority of *PpCLE1∷NGG*, *PpCLE8∷NGG* and *PpCLV1a∷NGG* filaments, all other lines accumulated signal in caulonemal tip cells. Stain was also detected in a subset of lines in other caulonemal cells (*PpCLE4∷NGG*, *PpCLE5∷NGG*, *PpCLE6∷NGG*, *PpCLE9∷NGG*, *PpCLV1b∷NGG*, *PpRPK2∷NGG*), in new branch initials (*PpCLE5∷NGG*, *PpCLE7∷NGG*, *PpCLE9∷NGG*, *PpCLV1b∷NGG*, *PpRPK2∷NGG*) or in chloronemal branch cells (*PpCLE4∷NGG*, *PpCLE9∷NGG*). Both *PpCLV1b∷NGG* and *PpRPK2∷NGG* accumulated stain in caulonemal tip cells and new branch/bud initials but *PpRPK2∷NGG* stained more strongly in caulonemal tip cells, and a subset of samples showed only this signal (4/37), while a subset of *PpCLV1B∷NGG* samples accumulated signal only in the branch initials (12/56). Numbers in each panel indicate the proportion of filaments displaying a similar expression pattern. Tissue was stained for 7.5 h (*PpCLE3∷NGG*, *PpCLE4∷NGG*, *PpCLE5∷NGG*, *PpCLE6∷NGG*, *PpCLE8∷NGG*, *PpCLV1b∷NGG*, and *PpRPK2∷NGG*), 15 h (*PpCLE2∷NGG*, *PpCLE9∷NGG* and *PpCLV1a∷NGG*) or 21 h (*PpCLE1∷NGG* and *PpCLE7∷NGG*). Black arrowheads indicate the strongest expression, grey arrows indicate weaker expression. Purple frames indicate lines with strong expression in caulonemal tip cells. Scale bars = 200 μm.

To observe expression at later developmental stages, we dissected caulonemal filaments extending from plants’ foraging fringe (Figure 1D). Whilst no signal was detected in *PpCLE1∷NGG*, *PpCLE8∷NGG* or *PpCLV1a∷NGG* lines, the promoters of all remaining *PpCLEs* were active in caulonemal tip cells, and *PpCLE5∷NGG* and *PpCLE9∷NGG* lines exhibited intense staining with a gradient of expression that tapered from the tip and extended to lateral initials. *PpCLV1b∷NGG* and *PpRPK2∷NGG* expression was detected in overlapping domains: whilst the *PpCLV1b* promoter was more active in the second and third cells from the tip and in early branch initials, *PpRPK2* promoter activity was reported preferentially in caulonemal tip cells. Taken together these expression data indicate roles for *PpCLE5*, *PpCLE9*, and *PpRPK2* in CLE production and perception in caulonemal tip cells, and to a lesser extent indicate potential roles *PpCLV1b* in caulonemal tip cells and branch initials.

### *clavata* mutants have defects in plant spread, plant shape and cell identity and *Pprpk2* mutants are larger with more irregular shapes than wild-type plants

To investigate roles for CLAVATA in protonemal development, we first quantified overall plant spread in WT and mutant plants (Figure 2A, B). We also quantified a measure of plant shape, the perimeter ratio. This reflects the circularity of plant spread such that a value of 1 corresponds to a perfectly circular shape, and higher values indicate an increase in the perimeter without an increase in the area (Figure 2B). Irregular plant shapes with high perimeter ratios indicate increased caulonema production. Whilst *PpcleAmiR1-3* mutants had a similar size and shape to wild-type plants, *PpcleAmiR4-7* mutants had greater spread but similar perimeter ratios to wild-type plants, implying increased but uniform protonemal growth (Figure 2B). Amongst receptor mutants, *Ppclv1a*, *Ppclv1b* and *Ppclv1a1b* mutants sometimes had increased areas and perimeter ratios, but the size and shape differences from wild-type plants were subtle and variable between experimental replicates. In contrast, *Pprpk2* and *Ppclv1a1brpk2* mutants were significantly larger with higher perimeter ratios than wild-type plants, indicating higher caulonema production and a key role for *PpRPK2* in plant size and shape determination (Figure 2B).

**Figure 2:**
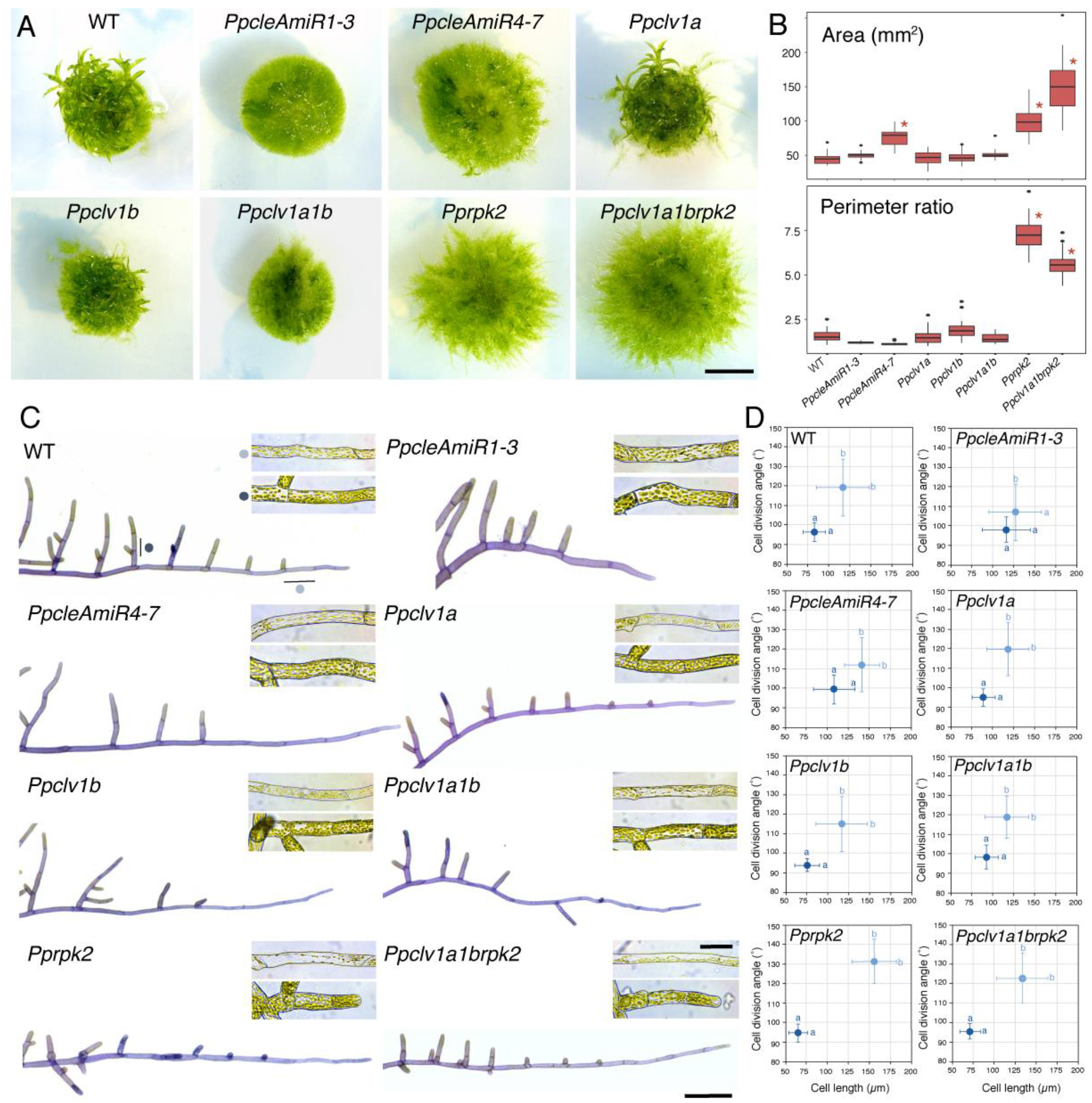
CLAVATA pathway components regulate plant shape and protonemal identity. **(A)** Images of four week-old plants grown in continuous light showing overall morphology. Scalebar = 5 mm. **(B)** Plant spread in *PpcleAmiR4-7*, *Pprpk2* and *Ppclv1a1brpk2* mutants was greater than wild-type plant spread, and triple mutants were larger than *Pprpk2* mutants. *Pprpk2* and *Ppclv1a1brpk2* mutants had a higher perimeter ratio than wild-type plants. Data from one of three experimental replicates are shown (n = 32). Asterisks indicate significant difference from wild-type (One-way ANOVA and Tukey test; p < 0.05). **(C)** Light micrographs of toluidine stained and dissected protonemal filaments, showing differences in apical dominance and cell morphology between genetic backgrounds. Insets show typical unstained subapical and branch cells for each line. Main scalebar = 200 μm, inset scalebar = 50 μm. **(D)** Subapical cells (dark blue dots; caulonemal cells in wild-type) and branch cells (pale blue dots; chloronemal cells in wild-type) have distinct shapes in wild-type plants, *PpcleAmiR4-7, Ppclv1a*, *Ppclv1b*, *Ppclv1a1b*, *Pprpk2* and *Ppclv1a1brpk2* mutants, but *PpcleAmiR1-3* mutant cell types are less distinct. Subapical cell and branch cell measurements partially overlap in all lines except *Pprpk2*. Different letters indicate significant difference in cell length or division plane orientation, bars indicate standard deviation (n ≥ 28; Two-way ANOVA p < 0.05).

To further investigate the cellular basis of plant spread and perimeter ratio differences between lines, differences in protonemal morphology were quantified by measuring the length and cell division plane orientations of cells in main filaments and side branches. These measures report differences in cell identity, and subapical cells in caulonemata (subapical cells) are typically longer with more oblique cell division plane orientations than cells in chloronemal branches (branch cells) [28] (Figure 2C, 2D). *Ppclv1a*, *Ppclv1b* and *Ppclv1a1b* mutants showed no significant differences from wild-type plants with respect to subapical and branch cell length or cell division plane orientation in most of the experimental replicates performed (Figure S3). However, although *PpcleAmiR1-3* mutants were the same size as wild-type plants, their cell types showed a mix of chloronemal and caulonemal characteristics, with less oblique cell divisions in subapical cells, and longer branch cells than wild-type plants (Figure S3). *PpcleAmiR4-7* mutant protonemata had longer subapical and branch cells than wild-type plants but retained distinct cell identities. Cell identity distinctions were enhanced in *Pprpk2* mutants, and sometimes in *Ppclv1a1brpk2* mutants which had subapical cells that were longer and with more oblique cell division plane orientations than wild-type cells (Figure S3).

More subtle qualitative differences in protonemal morphology were also observed following dissection of chloronemata and caulonemata and staining with toluidine blue (Figure 2C). Wild-type filaments initiate branches from the second subapical cell, and their continued growth gives the branching pattern a regular comb-like appearance, with teeth that are progressively larger moving away from the tip (Figure 2C). There was variation in the relative growth of main and branch filaments, such that branches more proximal to the tip were longer in *PpcleAmiR1-3* mutants than wild-type branch filaments in equivalent positions (Figure 2C). Conversely, side branches were shorter in *Pprpk2* and *Ppclv1a1brpk2* mutants than in wild-type plants compared to the main filament. Overall, expression and mutant phenotype analyses suggest that the CLAVATA pathway regulates protonemal morphology in *P. patens* and that *PpRPK2* holds the main role as a repressor of plant spread, perimeter ratio and caulonemal identity. As *Ppclv1a* and *Ppclv1b* mutant phenotypes were not fully penetrant, and *PpcleAmiR* mutant phenotypes reflect changes in expression of several *PpCLEs*, we focused further functional analyses on *Ppclv1a1b*, *Pprpk2* and *Ppclv1a1brpk2*.

### *Pprpk2* mutants are hypersensitive to cytokinin application

In wild-type plants, filament identity reflects a hormonal interplay between auxin and cytokinin, and cytokinin suppresses caulonemal identity [17]. We therefore reasoned that *PpRPK2* could promote cytokinin biosynthesis or enhance plants’ response to cytokinin to regulate plant spread and shape. To test the hypothesis that PpRPK2 promotes cytokinin biosynthesis we used LC-MS/MS to quantify cytokinin levels in protonemal homogenates of *Ppclv1a1b*, *Pprpk2* and *Ppclv1a1brpk2* mutant lines and wild-type controls. In flowering plants, cytokinin bases, and to a lesser extent ribosides are the active forms, while nucleotides and *O*-glucosides (OG) are inactive and can function in storage [29]. In *P. patens* isopentenyl adenine (iP), *trans*-zeatin (*t*Z) and their corresponding ribosides were shown to have the highest biological activity in a bud induction assay [30], while *cis*-zeatin (*c*Z), *cis*-zeatin riboside (*c*ZR) and all the ribotides had no bud inductive role. Twenty-six types of cytokinin were assayed, including free bases as iP, *t*Z, *c*Z, and dihydrozeatin (DHZ) and their ribotide (MP), riboside (R), and glycoside derivatives (Figure S4). Thirteen types of cytokinins were detected, and while some mutant-specific differences were present in the level of conjugates like *c*ZRMP, *t*ZOG and *t*ZROG, the only free base present at a higher concentration in the mutants was *c*Z in *Ppclv1a1b* mutants. No difference in the overall level of cytokinin was detected. Since no reduction in bioactive cytokinin was detected in *clavata* mutants, the hypothesis that PpRPK2 reduces plant spread by promoting cytokinin biosynthesis was rejected.

We next tested the hypothesis that PpRPK2 enhances the response to cytokinin by growing mutants and wild-type plants on media containing a solvent control (0.07% EtOH) or 100 nM of the synthetic cytokinin 6-benzylaminopurine (BAP), and analyzed their phenotypes after 4-5 weeks of growth (Figure 3). As expected, wild-type plants showed a decrease in plant spread in response to exogenously applied BAP (Figure 3A, B). On average the area of wild-type plants grown on 100 nM BAP was 26 ±11 % smaller than control plants, with no difference in perimeter ratio. The response to BAP treatment was sometimes dampened in *PpcleAmiR1-3* and *Ppclv1a1b* mutants, but *Pprpk2* and *Ppclv1a1brpk2* plants grown on 100 nM BAP consistently showed an enhanced response, with respectively a 59 ± 3% and 43 ± 9% decrease in plant spread and a reduction in perimeter ratio (Figure 3A, B).

**Figure 3:**
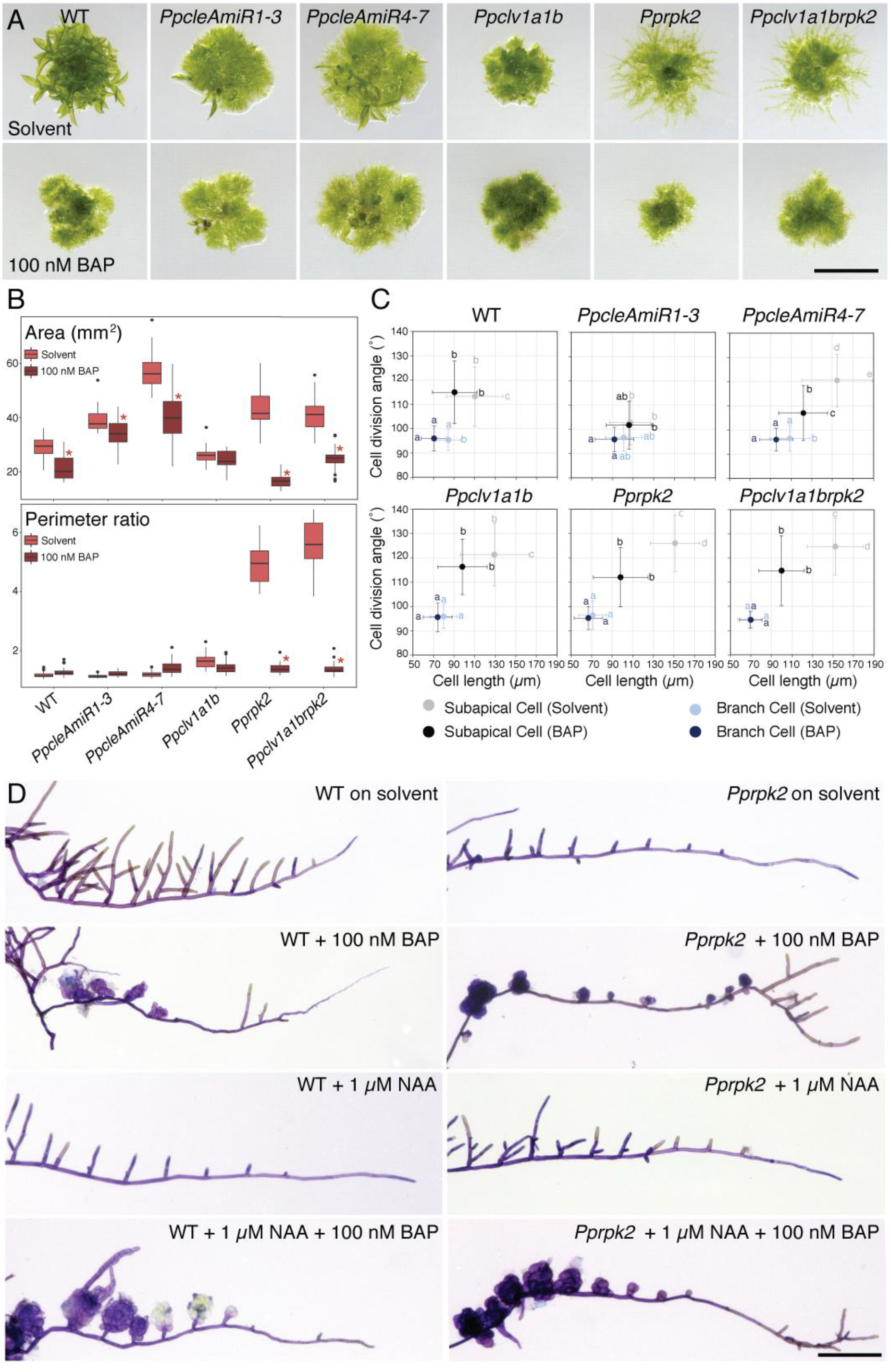
Cytokinin treatment disproportionally affects protonemal growth in clavata mutants. **(A)** Micrographs of four week-old plants grown on media containing a solvent control (EtOH) or cytokinin (100 nM BAP). Scale bar = 5 mm. **(B)** Graphs showing the area and perimeter ratio of solvent control and cytokinin treated plants. Whilst *Ppclv1a1b* plant area was not significantly affected by cytokinin treatment, wild-type, *PpcleAmiR1-3*, *PpcleAmiR4-7*, *Pprpk2* and *Ppclv1a1brpk2* plants show decreased area, with the decrease being more conspicuous in *Pprpk2* and *Ppclv1a1brpk2* mutants. The perimeter ratio did not change in response to cytokinin treatment in wild-type, *PpcleAmiR1-3*, *PpcleAmiR4-7* or *Ppclv1a1b* plants, but strongly decreased in *Pprpk2* and *Ppclv1a1brpk2* mutants. Asterisks indicate significant difference to EtOH control (Multi-way ANOVA and Tukey’s test; p < 0.05; n ≥ 30). **(C)** Graphs showing a reduction in subapical cell length in response to cytokinin application in wild-type and *Ppclv1a1b* plants, and a strong reduction in subapical cell length and division plane angles in *PpcleAmiR4-7*, *Pprpk2* and *Ppclv1a1brpk2* mutants grown on 100 nM BAP. Letters indicate significant differences between groups, error bars indicate standard deviation (n ≥ 90; Multi-way ANOVA and Tukey’s test; p < 0.05). **(D)** Micrographs of caulonemal filaments dissected from wild-type or *Pprpk2* mutants grown on plates containing a solvent control (EtOH), 100 nM BAP, 1 μM NAA, or 1 μM NAA + 100 nM BAP. *Pprpk2* mutants treated with BAP produced a near constitutive overbudding phenotype normally seen in wild-type plants treated with 1 μM NAA and 100 nM BAP. Scale bar = 200 μm.

To study responses to cytokinin at the cellular level, we dissected foraging filaments from plants and measured cell lengths and division planes as previously described (Figure 3C). In wild-type plants, BAP treatment reduced cell lengths in both subapical and branch cells, while cell angles were unaffected. In *Ppclv1a1b* and *PpcleAmiR1-3* mutant subapical cells respectively, responses to cytokinin were similar or weaker than wild-type responses. However, *PpcleAmiR4-7*, *Pprpk2* and *Ppclv1a1brpk2* subapical cells showed a large reduction in both cell length and division plane angle following BAP treatment. Thus, *Pprpk2* and *Ppclv1a1brpk2* mutant plants have an enhanced response to cytokinin in caulonemal tip cells as well as in whole plants, refuting the hypothesis that *PpRPK2* promotes the cytokinin response. *Pprpk2* mutants showed further evidence of cytokinin hypersensitivity in gametophore initiation, which is normally upregulated by cytokinin [17]. Whilst wild-type plants grown on BAP showed a higher frequency of gametophore initiation than untreated controls (Figure 3D), *Pprpk2* mutants grown on BAP showed almost constitutive gametophore initiation. In wild-type plants, such bud production is induced following combined BAP and NAA treatment (Figure 3D) [31].

### Auxin synthesis is not elevated in *Pprpk2* mutants, but mutant increases in plant spread require auxin

Because auxin can enhance or suppress cytokinin activity in *P. patens* [17], we hypothesized that cytokinin hyper-sensitivity in *Pprpk2* and *Ppclv1a1brpk2* mutants could reflect perturbations in auxin homeostasis. To test whether *clavata* mutant phenotypes depend on auxin biosynthesis, we grew wild-type and mutant plants on media containing either a pharmacological inhibitor of auxin synthesis (10 μM L-Kynurenine) or a solvent control (0.01 % DMSO) (Figure 4A–C). When grown on L-Kyn, wild-type plants showed a significant decrease in area relative to plants grown on a solvent control, and *clavata* mutant plants showed a similar decrease in area to wild-type plants, but *PpcleAmiR4-7*, *Pprpk2* and *Ppclv1a1brpk2* plants showed a larger decrease (Figure 4B). Perimeter ratios were invariant between treatments in wild-type, *PpcleAmiR1-3*, *PpcleAmiR4-7* and *Ppclv1a1b* plants, but decreased significantly in *Pprpk2* and *Ppclv1a1brpk2* plants grown on 10 μM L-Kyn, indicating that the irregular shapes of these mutants depend on auxin synthesis. To assay the effects of L-Kyn on cell identity, we dissected foraging filaments and measured cell length and cell division plane orientations as previously described. While no response to L-Kyn in branch cells was detected in any genotype, subapical cells of all lines were shorter and/or had a division plane angle closer to 90° when plants were grown on L-Kyn, and these differences were significant in *PpcleAmiR4-7*, *Ppclv1a1b* and *Pprpk2* mutants. Hence, wild-type and *clavata* mutant plants have a qualitatively similar response to auxin biosynthesis inhibition, which reduces the morphological distinction between subapical and branch cells. These data indicate that active auxin synthesis is needed for the caulonemal overproliferation phenotype of *Pprpk2* and *Ppclv1a1brpk2* mutants.

**Figure 4:**
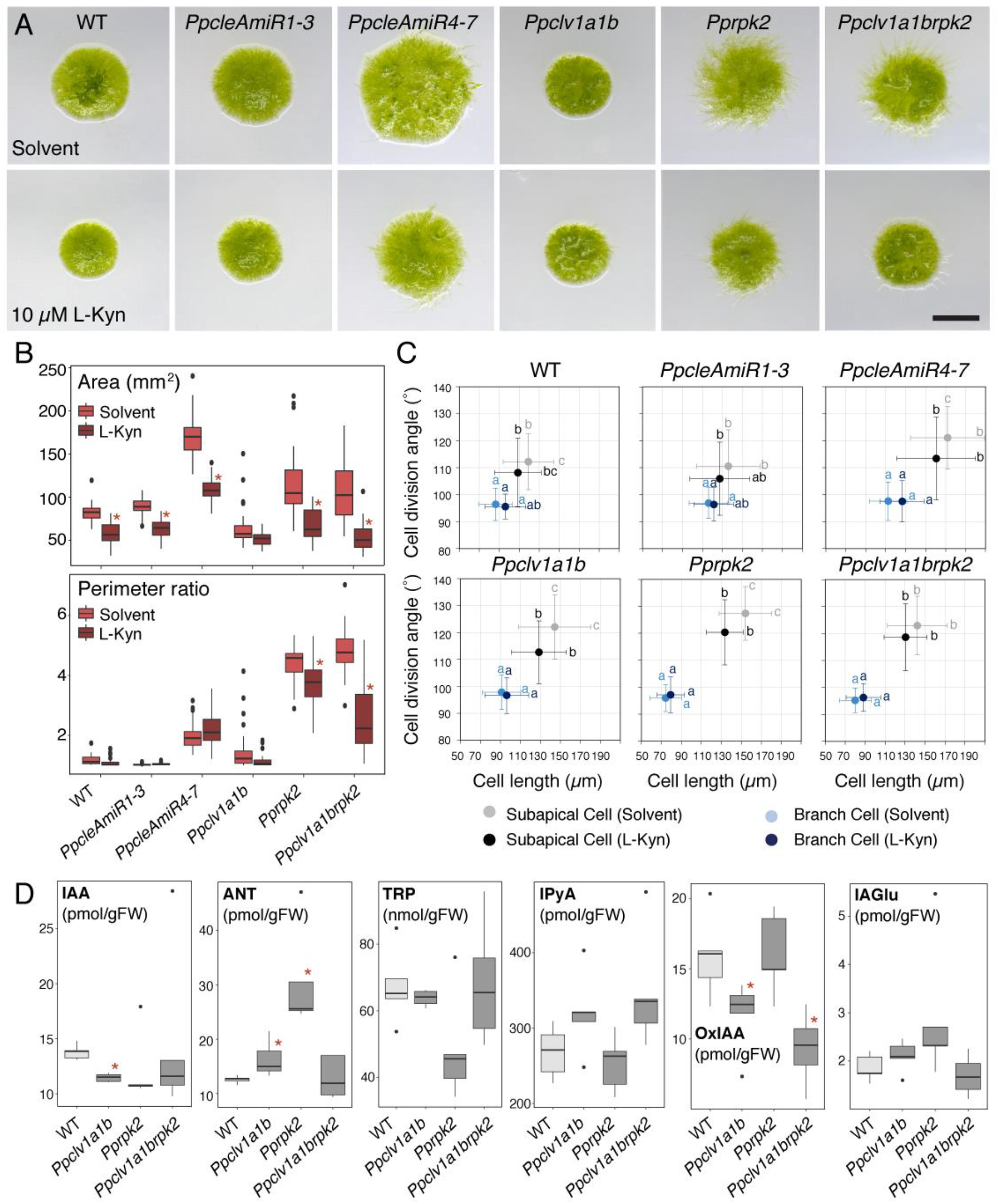
*Pprpk2* and *Ppclv1a1brpk2* mutant phenotypes are auxin synthesis dependent, and mutants have synthesis defects. **(A)** Micrographs of four week-old plants grown on media containing a solvent control (DMSO) or a pharmacological inhibitor of auxin synthesis (10 μM L-Kyn). Scale bar = 5 mm. **(B)** Plant spread in all lines diminishes in response to L-Kyn, but perimeter ratio values decreased only in *Pprpk2* and *Ppclv1a1brpk2* mutants (Two-way ANOVA and Tukey’s test, n ≥ 30, * = p value < 0.05 between treatment and control). **(C)** L-Kyn treatment suppresses subapical cell length in *Ppclv1a1b* and *Pprpk2* mutants, and suppresses oblique cell divisions in *PpAmiR4-7*, *Ppclv1a1b* and *Pprpk2* mutants. Letters indicate significant differences between groups, error bars indicate standard deviation (Multi-way ANOVA and Tukey’s test, n ≥ 90, p value < 0.05). **(D)** Quantification of auxin metabolites from wild-type, *Ppclv1a1b*, *Pprpk2* and *Ppclv1a1brpk2* protonemal cultures showed that auxin levels were depleted in *Ppclv1a1b* and *Pprpk2* mutants in 4/5 biological replicates. IAA = Indole-3-acetic acid, ANT = Anthranilate, TRP = Tryptophan, IPyA = Indole-3-pyruvic acid, OxIAA = 2-oxindole-3-acetic acid, IAGlu = IAA-Glucose; n = 5, * = p value <0.05.

As auxin promotes caulonemal development, we hypothesised that auxin overproduction could account for *Pprpk2* and *Ppclv1a1brpk2* mutant phenotypes. We therefore quantified biologically active auxin (indole-3-acetic acid, IAA), its precursors (anthranilate, ANT; L-tryptophan, TRP; and indole-3-pyruvic acid, IPyA) and degradation products (IAA-glutamate, IAGlu; and 2-oxindole-3-acetic acid, OxIAA) from protonemal homogenates of wild-type and mutant plants. However, we found that *Ppclv1a1b* and four out of five replicates of *Pprpk2* mutant tissue batches in fact had lower IAA content than wild-type protonemata, while *Ppclv1a1brpk2* IAA levels were variable (Figure 4D). *Ppclv1a1b* and *Pprpk2* protonemata contained more ANT, than wild-type protonemata, and *Ppclv1a1b* and *Ppclv1a1brpk2* mutant protonemata contained less OxIAA than wild-type samples. Thus, *Ppclv1a1b* (and usually *Pprpk2* mutant) protonemata contained less biologically active auxin (IAA) than wild-type protonemata, and we rejected our hypothesis that upregulation of auxin synthesis contributes to *Pprpk2* and *Ppclv1a1brpk2* mutant phenotypes.

### *Pprpk2* mutants manifest a perturbed developmental response to auxin

Enhanced sensitivity to auxin could yield similar developmental outcomes to elevated auxin levels, and to evaluate the response of *clavata* mutants to exogenous auxin, we grew plants on media containing either the synthetic auxin 1-naphthaleneacetic acid (1 μM NAA) or a solvent control (0.07 % EtOH) (Figure 5A–C). In line with previous studies [17, 21], wild-type plants grown with additional auxin showed increased areas relative to plants grown on a solvent control, but wild-type perimeter ratios were not affected. While *PpcleAmiR1-3*, *PpcleAmiR4-7* and *Ppclv1a1b* plants showed the same response as wild-type plants, *Pprpk2* plants showed an opposite response, with a decrease in both area and perimeter ratio (Figure 5A–B). At the cellular level, only subapical cells showed a strong response to NAA application, attaining division angles closer to 90° in wild-type plants. *PpcleAmiR4-7* and *Ppclv1a1brpk2* mutants also showed significant reductions in division angle following auxin application, and *Ppclv1a1b* and *Pprpk2* mutants showed significant reduction in subapical cell lengths. Thus, auxin, L-Kyn and cytokinin all affected caulonemal morphology by modulating subapical cell but not branch cell shape in wild-type and mutant plants. The longer and more oblique division subapical cell phenotypes of *Pprpk2* mutant caulonemata were never replicated in wild-type plants.

**Figure 5:**
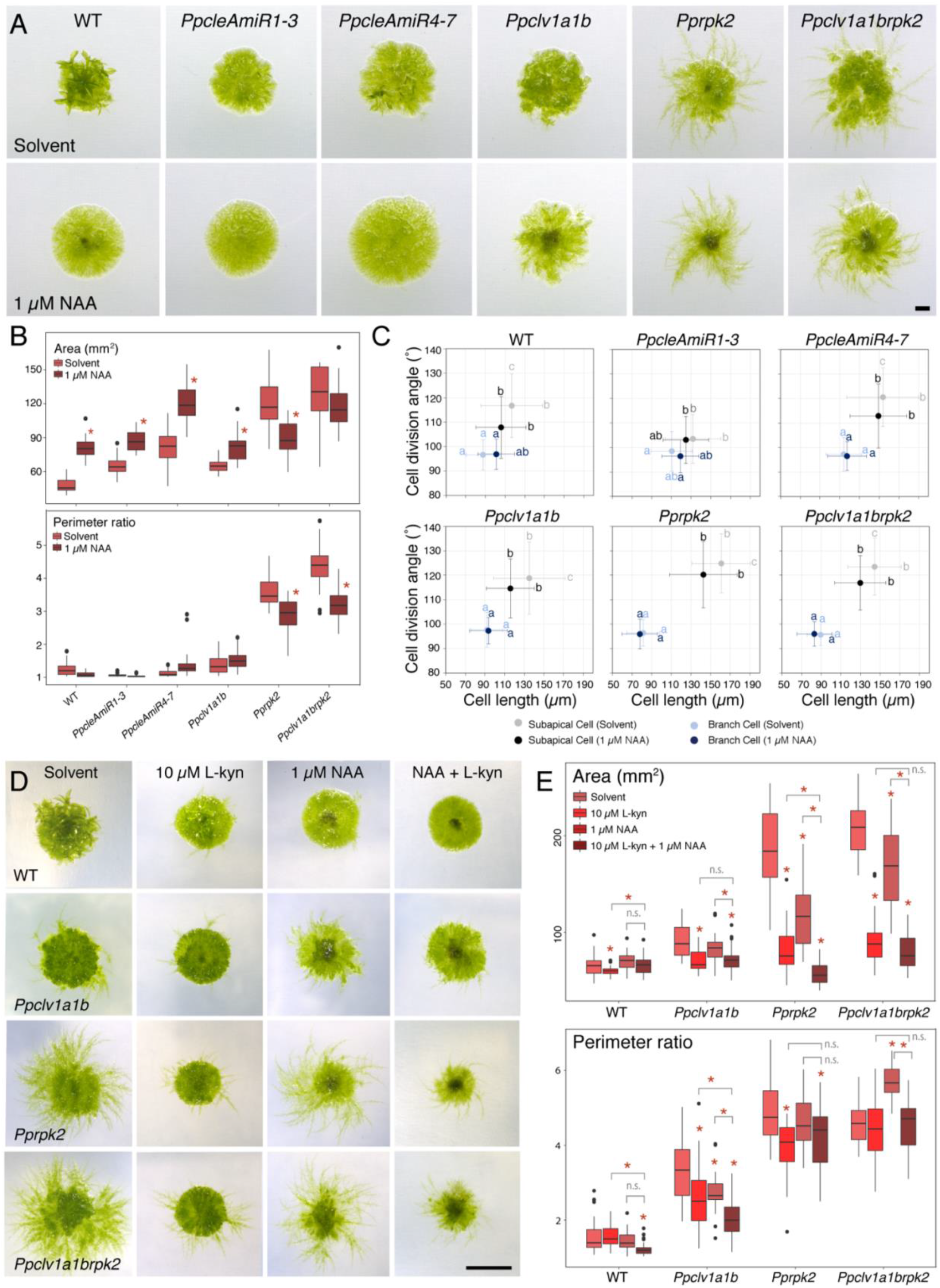
*Pprpk2* and *Ppclv1a1brpk2* mutants manifest an abnormal auxin response. (A) Micrographs of four week-old plants grown on media containing a solvent control (EtOH) or auxin (1 μM NAA). Scale bar = 1 mm. (B) Quantitative analyses showed that *Pprpk2* and *Ppclv1a1brpk2* mutant plants show an auxin-dependent decrease in plant spread and perimeter ratio, whilst all other backgrounds increased in area in response to auxin treatment and showed no change in perimeter ratio (Two-way ANOVA and Tukey’s test. n ≥ 30. * = p value < 0.05 between treatment and control). (C) Wild-type, *PpcleAmiR4-7* and *Ppclv1a1brpk2* subapical cell division angles diminished following 1 μM NAA treatment, and *Ppclv1a1b* and *Pprpk2* subapical cell lengths decreased, but branch cells and *PpcleAmiR1-3* mutant cells showed no change in length or division angle. Letters indicate significant differences between groups, error bars indicate standard deviation (Multi-way ANOVA and Tukey’s test; n ≥ 90 for all other genotypes; p value < 0.05). (D) Micrographs of four week-old plants grown on media containing a solvent control (0.07 % EtOH + 0.01 % DMSO), 10 μM L-Kynurenine (L-Kyn), 1 μM NAA or a combination of 10 μM L-Kyn and 1 μM NAA. Scalebar = 5 mm. **(E)** Quantitative analyses showed that whilst wild-type plant spread showed little response to exogenously applied auxin (1 μM NAA), auxin synthesis inhibitors (10 μM L-Kyn) or a combination of 1 μM NAA and 10 μM L-Kyn, *Pprpk2* mutant spread strongly decreased in all treatments and the combined treatment led to the strongest decrease. The response of *Ppclv1a1b* and *Ppclv1a1brpk2* mutants varied between experimental replicates, as did perimeter ratios. Asterisks above data indicate significant difference from untreated controls, asterisks above bars indicate significant difference between treatments of interest (n ≥ 30; One-way ANOVA and Tukey test on each genotype; p < 0.05).

To further dissect the effects of auxin synthesis and auxin response, we uncoupled these two processes by growing plants on media containing either a solvent control (0.01 % DMSO + 0.07 % EtOH), 10 μM L-Kyn (reduced synthesis), 1 μM NAA (response plus endogenous synthesis), or a combination of 10 μM L-Kyn and 1 μM NAA (response in absence of synthesis). We reasoned that should mutant phenotypes be caused by enhanced auxin perception or response, mutants would respond similarly to saturating concentrations of exogenously applied auxin both with (1 μM NAA treatment) and without endogenous synthesis (combined treatment). Wild-type plant spread increased in L-Kyn + NAA treated plants relative to L-Kyn treated plants, illustrating the normal response to exogenously applied auxin in the absence of auxin synthesis (Figure 5D,E). Whilst responses to exogenous auxin in *Ppclv1a1b* and *Ppclv1a1brpk2* mutants were somewhat variable between experimental replicates, *Pprpk2* mutants grown on the combined treatment had reduced areas relative to either single treatment (Figure 5D,E). These data did not fit our hypothesis that *Pprpk2* mutants should respond similarly to exogenously applied auxin regardless of the level of endogenous auxin synthesis, and thus our hypothesis was insufficient to fully account for the phenotype of *Pprpk2* mutants.

### *Pprpk2* mutants manifest auxin transport perturbations

The L-Kyn + NAA treatment above would not only perturb the amount of auxin present, but also its spatial distribution, so we next hypothesized that local auxin gradients could be an important shape determinant in *Pprpk2* mutants. Promoter fusions have shown that *P. patens PIN* genes are highly expressed in protonemal tip cells, and the chloronemal to caulonemal transition is accelerated in *pinapinb* mutants but suppressed in *PINA* and *PINB* (and to a lesser extent *PINC*) overexpressors [20]. To investigate a potential contribution of auxin transport to the *Pprpk2* mutant phenotype, we first used a pharmacological approach to assay the sensitivity of mutants to transport inhibition, growing wild-type, *Ppclv1a1b*, *Pprpk2* and *Ppclv1a1brpk2* mutants on media containing a solvent control, 5 μM of the auxin transport inhibitor naphthylphthalamic acid (NPA) [32], 1 μM NAA or a combination of 5 μM NPA and 1 μM NAA (Figure 6 A,B). Wild-type plant spread increased following treatment with auxin in both the single and combined treatment, but perimeter ratios decreased following NPA treatment, in both the single and combined treatment, consistent with the notion of a more uniform and widespread chloronema to caulonema transition than usual. Whilst *Ppclv1a1b* mutant shape showed a similar response to wild-type plants to pharmacological agents, *Pprpk2* and *Ppclv1a1brpk2* mutants showed mild growth suppression following treatment with 5 μM NPA or 1 μM NAA, and strong growth suppression following the combined treatment, consistent with the auxin overaccumulation phenotypes shown in Figure 4. We therefore concluded that *Pprpk2* and *Ppclv1a1brpk2* mutants were hypersensitive to auxin transport inhibition. To assay the molecular basis of sensitivity to auxin transport inhibition, we quantified *PIN* expression in protonemal homogenates of wild-type and *clavata* plants by qPCR (Figure 6C). Whilst *PIND* expression showed no variation between wild-type plants and mutants, *PINA* expression was reduced in *Pprpk2* mutants and *PINB* expression was reduced in all mutants (Figure 6C). *PINC* is expressed at around the PCR detection limit in wild-type protonemal homogenates [33], and we were unable to detect expression in any mutant. Thus, suppression of canonical *PIN* expression is likely to contribute to generation of the *Pprpk2* mutant phenotype by altering the auxin distribution.

**Figure 6:**
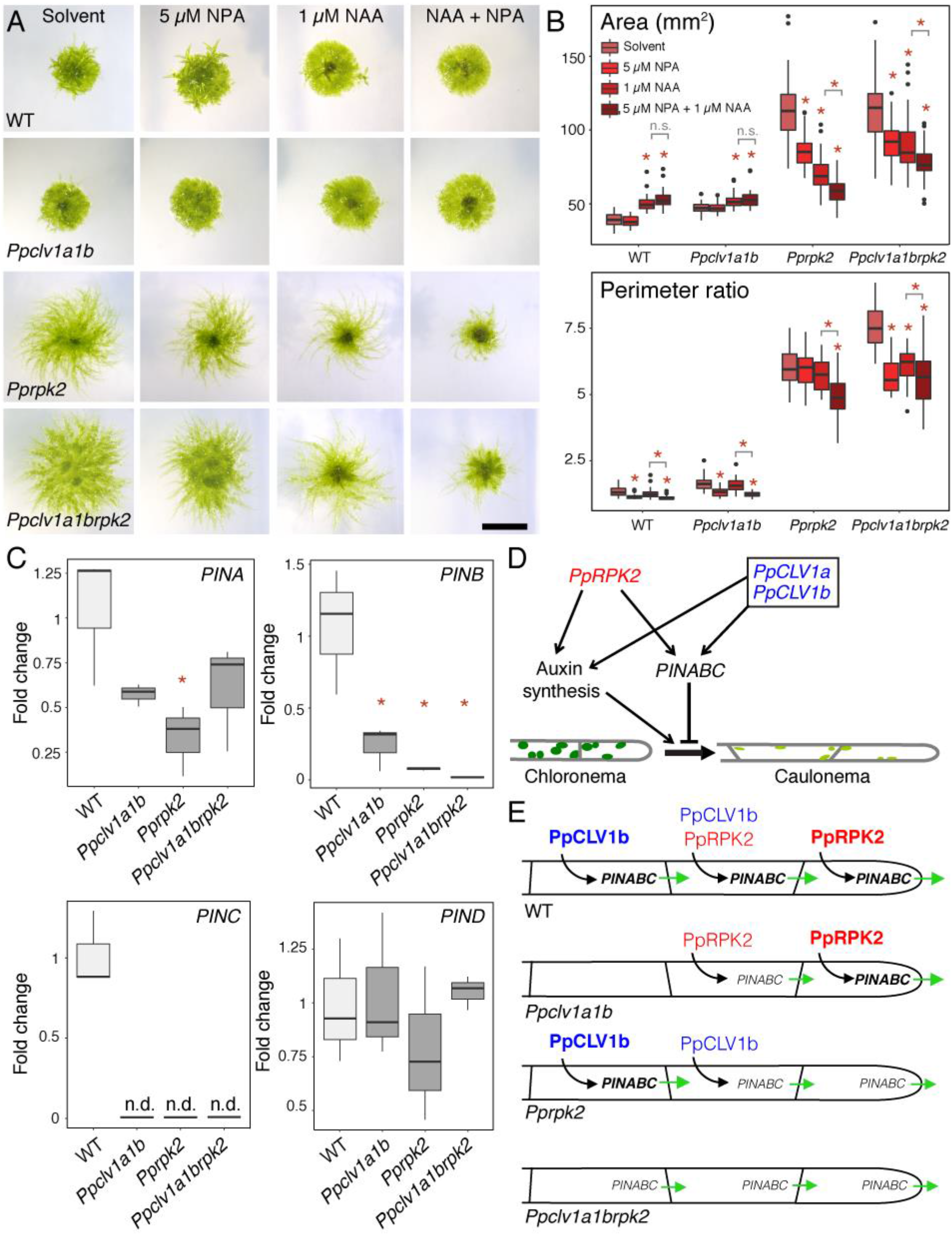
PIN-mediated auxin transport is dampened in *clavata* mutants. **(A)** Images of four week-old plants grown on a solvent control, 5 μM NPA, 1 μM NAA or a combination of 5 μM NPA and 1 μM NAA. Scalebar = 5 mm. **(B)** Quantitative analyses showed that *Pprpk2* and *Ppclv1a1brpk2* mutant plants showed auxin- and auxin transport-dependent decreases in plant spread and perimeter ratio, whereas wild-type and *Ppclv1a1b* plants showed increased plant spread and transport-dependent reductions in perimeter ratio. Asterisks above data indicate significant difference from untreated plants of the same genotype, asterisks above bars indicate significant difference between treatments of interest (n ≥ 30; One-way ANOVA and Tukey test on each genotype; p < 0.05). **(C)** qPCR showed that *PINA* expression was depressed in *Pprpk2* mutants relative to wild-type plants, and *PINB* and *PINC* expression were depressed in all mutants. *PIND* expression showed similar expression levels in wild-type plants and mutants. *60S* was used as housekeeping control (n = 3; n.d. = not determined; * = p value < 0.05 between wild-type and mutant samples. No significant differences were found between mutants. **(D)** Model of the CLAVATA gene regulatory network involved in protonemal stem cell identity. Black arrows indicate positive regulation, black T bar indicates negative regulation. **(E)** Model of CLAVATA function in wild-type and *clavata* mutant plants. Black arrows indicate positive regulation of gene expression, green arrows indicate the level and direction of auxin fluxes, and the size and strength of the font reflect the strength of protein of gene action.

## Discussion

The data discussed above led us to a model of CLAVATA function whereby CLAVATA modulates the activity of protonemal tip cells by promoting auxin biosynthesis and expression of *PINs A-C* and suppressing the chloronema to caulonema developmental transition to determine plant spread and shape (Figure 6D). As auxin synthesis promotes but transport represses caulonemal development [17, 20, 21], and CLAVATA promotes both synthesis and transport, our data point to a stronger role for auxin transport than synthesis in specifying caulonemal identity. Relative to wild-type plants, *Pprpk2* and *Ppclv1a1b* mutants show similar reductions in auxin content and *PIN* expression (Figure 4D and Figure 6C), but *Pprpk2* mutants show a much more severe mutant phenotype (Figure 2). We hypothesize that the dominant role of *PpRPK2* over *PpCLV1a* and *PpCLV1b* reflects differences in protonemal receptor gene expression patterns. While *PpRPK2* is strongly expressed in the apical cell of caulonemal filaments, *PpCLV1b* expression is strongest in subapical cells and *PpCLV1a* expression is only detectable in sporelings (Figure 1 D, Figure 6E). We propose that PpRPK2 controls the auxin transport status of the tip cell, directly affecting plant growth (Figure 6E). Following this model, the auxin transport status of the tip cell would be similar in *Pprpk2* and *Ppclv1a1brpk2* mutants, accounting for the similar phenotypes of these mutants, whereas *Ppclv1a1b* mutants would have an altered auxin distribution in subapical cells.

Recent work has shown that although protonemal tips cells are the distal site of phenotype determination, auxin signalling is concentrated at the centre of plants (proximally) [22, 34, 35]. Gain-of-function *pACT∷PpPINGFP* mutants do not produce caulonemata, and *PIN*s are expressed most strongly in protonemal tip cells and a few subapical cells [20]. Our model for PpRPK2 function fits with these prior data by suggesting *PIN* expression levels as a primary determinant of protonemal morphology. Protonemal tip cells show a low capacity for TIR/AFB- mediated auxin sensing but can be highly responsive to auxin if repressor ARF (PpARFb) levels are low, and PpARFb depletion by tasiRNAs in a subset of filaments patterns caulonema initiation and protonemal branching at plants’ foraging fringe [23]. *PpCLE* and *PpRPK2* expression are enriched in caulonemal tip cells, and as CLE peptides are diffusible, PpRPK2 could act in parallel or in sequence with PpARFb to pattern caulonema initiation. As CLAVATA genes are not expressed in secondary chloronemata, we propose that differences in the auxin response between chloronemata and caulonemata in wild-type could reflect distinct domains of CLAVATA and *PpARFb* activity.

From an evolutionary perspective, our work suggests that roles for CLAVATA in modulating PIN, auxin response and stem cell identity are conserved within the land plants [36–38]. We propose that a transcription factor X acts as an intermediary between CLAVATA and auxin homeostasis. Work in Arabidopsis puts WUSCHEL at an equivalent position to X in the *CLAVATA* gene regulatory network [2], and WUSCHEL maintains low auxin status in the stem cells of the central meristem zone [15]. WUSCHEL belongs to the T3 clade of the *WOX* gene family, which arose from a gene duplication predating the origin of vascular plants. The ancestral complement of land plant *WOX*es comprised the T1 *WOX* clade and a T2 + T3 *WOX* precursor lineage, which was lost in bryophytes [15]. In combination with functional work in *P. patens* and *Marchantia polymorpha* showing that CLAVATA and *WOX13L* (T1 *WOX*) genes regulate dissimilar developmental processes or act independently, our data suggest that *X* is unlikely to be a *WOX* gene [26, 27], and as the TDIF/PXY module regulates ARF action independently from WOX genes [38], we propose that *X* could be an ARF. Thus, downstream components of the CLAVATA gene regulatory network have been remodelled during evolution. WUSCHEL could have been co-opted into the CLAVATA gene regulatory network in euphyllophytes, a T3 WUSCHEL precursor could have been co-opted in the last common ancestor of vascular plants, or alternatively CLAVATA could have operated with a T2 + T3 *WOX* precursor to regulate auxin homeostasis in the last common ancestor of land plants.

## Methods

### Plant growth

Gransden was used as the wild-type strain for all experiments, and *PpcleAmiR1-3*, *PpcleAmiR4-7*, *Ppclv1a*, *Ppclv1b*, *Ppclv1a1b*, *Pprpk2*, *PpCLE1∷NGG*, *PpCLE2∷NGG*, *PpCLE7∷NGG*, *PpCLV1A∷NGG*, *PpCLV1B∷NGG* and *PpRPK2∷NGG* line generation strategies were previously described [25]. *Ppclv1a1brpk2* mutants were a gift from Joe Cammarata, Adrienne Roeder, and Mike Scanlon, and were generated by CRISPR-Cas9 editing *PpRPK2* in a *Ppclv1a1b* mutant background. Plants were spot-propagated on BCDAT media (0.5% Agar, 1 mM Magnesium sulphate (MgSO_4_), 3.67 mM monopotassium phosphate (KH_2_PO_4_), 10 mM potassium nitrate (KNO_3_), 45 μM iron sulphate (FeSO_4_), 5 mM ammonium tartrate dibasic ((NH_4_)_2_C_4_H_4_O_6_), 0.5 mM CaCl_2_, 1:1000 dilution of Trace Elements Solution (0.614mg/L H_3_BO_3_, 0.055mg/L AlK(SO_4_)_2_.12H_2_O, 0.055mg/L CuSO_4_.5H_2_O, 0.028mg/L KBr, 0.028mg/L LiCl, 0.389mg/L MnCl_2_.4H_2_O, 0.055mg/L CoCl_2_.6H_2_O, 0.055mg/L ZnSO_4_.7H_2_O, 0.028mg/LKI and 0.028mg/L SnCl_2_.2H_2_O)) unless otherwise stated, and grown at 23°C in continuous light or at 22°C in long day conditions (16h light/8h dark). To generate spores, protonemal homogenates were sown on peat plugs in Magenta vessels and grown at 23°C in continuous light for 8-10 weeks prior to transfer to 16°C short day conditions (16h dark/8h light). Mature sporogonia were harvested, incubated in 10 % sodium hypochlorite for 5 minutes and washed three times with sterile water, then refrigerated or ruptured in sterile water and germinated in continuous light on BCDAT medium lacking Trace Elements Solution and with 10 mM CaCl_2_.

### Pharmacological treatments

1 mM and 100 μM 1-naphthaleneacetic acid (NAA) and 100 μM and 10 μM 6-Benzylaminopurine (BAP) stock solutions were prepared in 70% ethanol. 100 mM stocks of L-Kynurenine (L-Kyn) were prepared using Dimethyl sulfoxide (DMSO) as a solvent. 5 mM N-1-naphthylphthalamic acid (NPA) stocks were prepared in 1 mL of DMSO made up to 50 mL with 70% EtOH. All reagents were added to warm growth media prior to pouring plates.

### Phenotype analysis

Plant areas and perimeters were measured using FIJI from images taken using a Keyence VHX-1000 microscope with a 10 X objective. These values were used to calculate the Perimeter Ratio, the ratio between the measured perimeter and the perimeter of a perfectly circular plant of the same area. For cell identity measurements, filaments protruding from the margins of four week-old plants were dissected and stained with 0.3 % Toluidine Blue for 2 minutes, rinsed in water and mounted on slides with coverslips prior to imaging with a Leica DM2000 microscope using a 40 X objective or a Keyence VHX-1000 microscope using a 50-200 X objective. The length and cell division angle of sub-apical cells of main filaments and of the second cell in branches with at least three cells were measured using FIJI as previously described [28].

### Generation of promoter∷NGG lines

Promoter sequences from *PpCLE3* (2427 bp), *PpCLE4* (2867 bp), *PpCLE5* (1731bp) and *PpCLE6* (1458 bp) were PCR amplified using a proofreading polymerase and cloned directly into the SmaI site of a modified PIG1NGGII [39] vector in which an NptII resistance cassette was substituted for the BSD cassette [25]. The promoter plus the first few amino acids of the peptide coding sequence were PCR amplified from *PpCLE8* (3216 bp) and *PpCLE9* (1799 bp) prior to insertion into PIG1NGGII to generate a translational fusion with the reporter gene. All constructs were linearised with PmeI prior to plant transformation. Lines were screened using a forward primer from the PIG1 targeting locus and a promoter-specific reverse screening primer to check the 5’ integration site, and a CaMV terminator forward primer and reverse primer from the *PIG1* locus were used to check the 3’ integration site. Southern analyses to verify targeting were undertaken using either a *GFP-GUS* probe PCR amplified to incorporate a DIG label (*PpCLE3*, *PpCLE4*, *PpCLE5*, *PpCLE6*, and *PpCLE9*) or a probe against the *35S∷NptII* resistance cassette (*PpCLE8*) as illustrated in Figure S1, and using methods described in Whitewoods et al. (2018) [25]. Primer sequences are listed in Table 1.

**Table 1:**
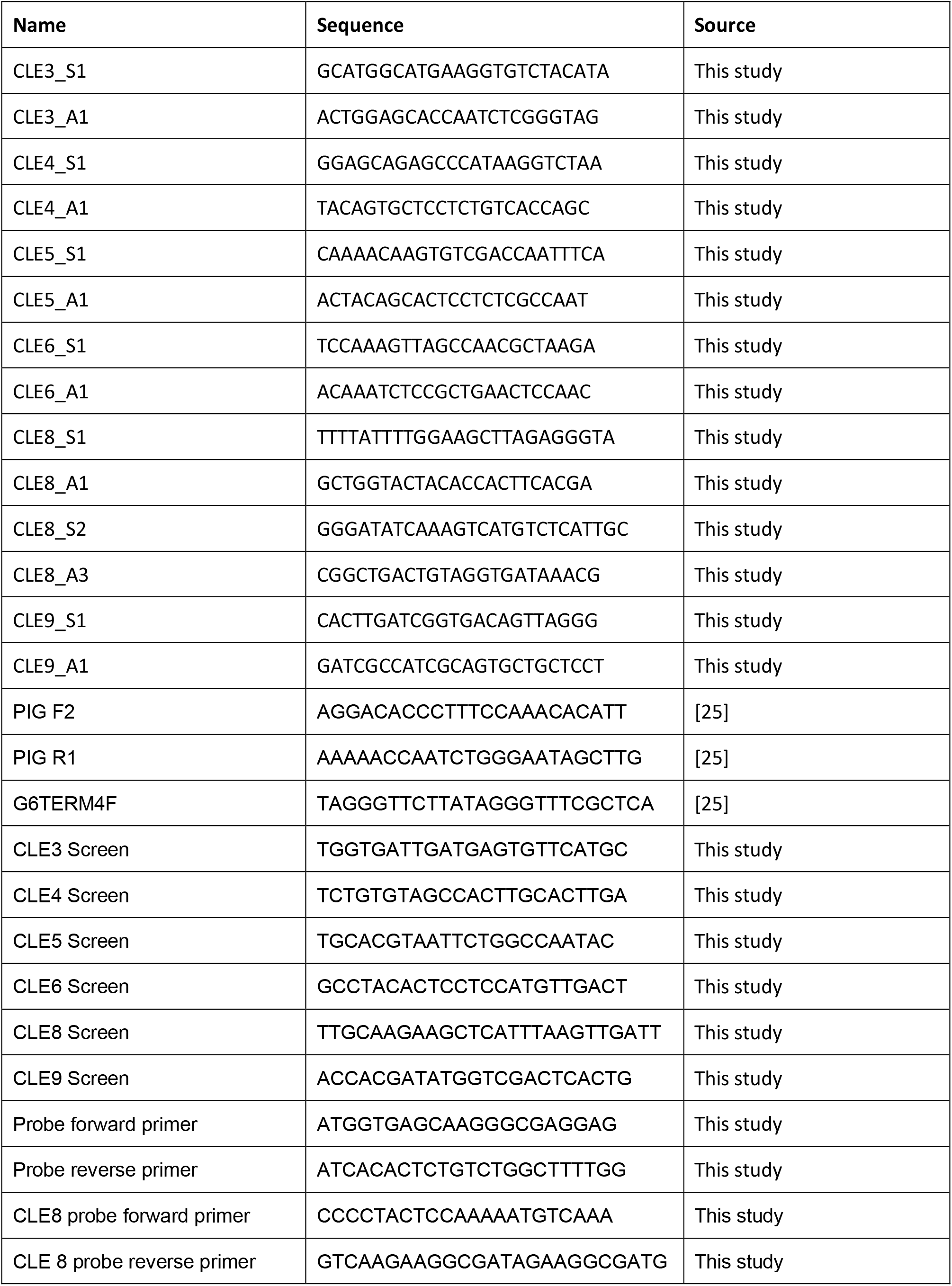
List of primers used for *Promoter∷NGG* line generation.

### Nucleic acid extraction

DNA for PCR and Southern analysis was extracted using a modified CTAB protocol [40] as described by [41]. RNA for expression analyses was extracted from 5 day-old protonemata using a RNeasy Plant Mini Kit (Qiagen). Genomic DNA removal and cDNA synthesis were performed with a Quantitect Reverse Transcription kit (Qiagen).

### Southern analysis

Southern blot analysis was carried out as described previously [42]. Genomic DNA (2.5 μg) was digested with *Hin*dIII (*PpCLE3*, *PpCLE4*, *PpCLE5*, *PpCLE8* and *PpCLE9*) or *Sca*I (*PpCLE6*), and transferred to HyBond-N-Plus membrane for hybridisation with the digoxygenin-labelled GFP-GUS reporter sequence.

### Expression analysis by qPCR

qPCR was performed using a SYBR Green Quantitect kit and a Stratagene Mx3005P bioanalyser with 95 °C 15 min, and then 94 °C for 15s, 60 °C for 30s, 72 °C 30scycling conditions for 40 cycles. Amplicon size was checked by dissociation curve. The efficiency of each primer pair was calculated on serial dilutions of wild-type cDNA, and only primer pairs with an efficiency between 90 % and 110 % were used in further experiments. To calculate fold change for each sample relative to wild-type expression levels, the ΔΔCt method was used [43]. *PINA*, *PINB*, *PINC*, *PIND* and *60S* genes were amplified using primers listed in Table 2.

**Table 2:**
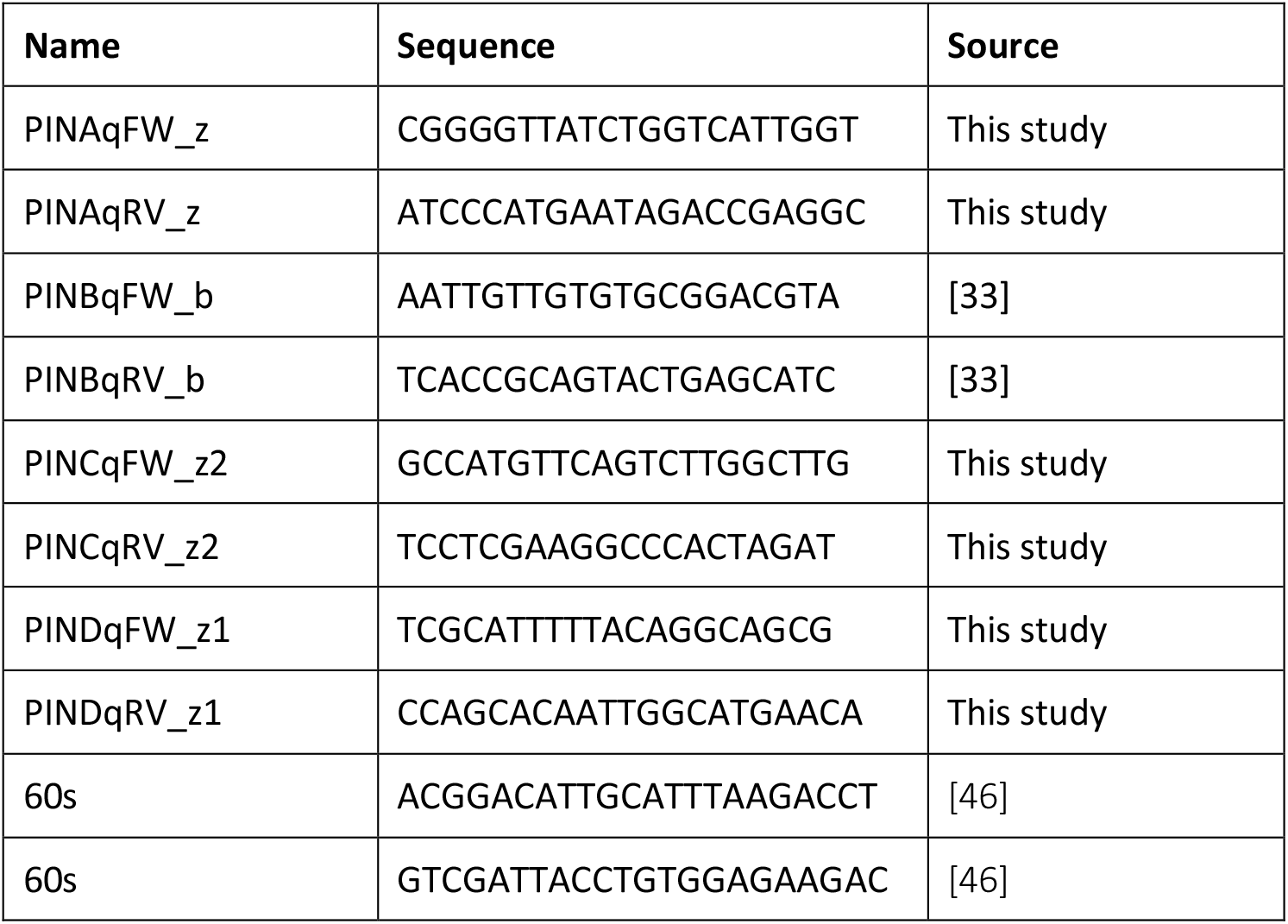
List of primers used for Q-PCR

### Histochemical expression analysis

For whole plant expression analyses, plants were grown on BCDAT media containing 0.5 % agar, cut out of plates and immersed in a staining solution which comprised 100 mM phosphate Buffer (pH 7.0), 10 mM Tris-HCl (pH 8.0), 1 mM ethylenediaminetetraacetic acid (EDTA), 0.05 % Triton X-100, 2 mM potassium ferricyanide, 2 mM potassium ferrocyanide and 1 mg/mL X-Gluc (5-Bromo-4-chloro-3-indolyl-ß-D-glucuronic acid) dissolved in 10 % (v/v) DMSO). Samples were incubated at 37 °C for 7.5 h (*PpCLE3∷NGG*, *PpCLE4∷NGG*, *PpCLE5∷NGG*, *PpCLE6∷NGG*, *PpCLV1b∷NGG* and *PpRPK2∷NGG*), 15 h (*PpCLE2∷NGG*, PpCLE8∷NGG, *PpCLE9∷NGG* and *PpCLV1a∷NGG*) or 21 h (*PpCLE1∷NGG* and *PpCLE7∷NGG*) except for samples in Figure 1A, which were incubated for a third of the time with 0.5 mM potassium ferricyanide and 0.5 mM potassium ferrocyanide instead. Reactions were stopped by substituting the staining solution with ethanol. Samples were bleached overnight in 70% ethanol, rinsed in water then imaged using a Keyence VHX-1000 digital microscope (whole plants) or a Leica DM2000 microscope (sporelings and filaments). For spore and sporeling expression analyses, plants were germinated on cellophane discs, moved to staining solution and imaged immediately after incubation.

### Hormone quantification

Hormone quantification by an ultra-high performance liquid chromatography-electrospray tandem mass spectrometry (UHPLC-MS/MS) was undertaken using protonemal homogenates grown through three passages of five days in continuous light. 50 mg of tissue was snap frozen in liquid nitrogen and hormone extraction and quantification was undertaken as described by Novák et al (2012) and Svačinová et al (2012) [44 and 45].

## Author contributions

Z.N.V. and J.H. conceived the project and designed the experiments. A.C. and Y.K. engineered *promoter∷NGG* fusion lines as part of the Leeds Moss Transformation Service. O.N. performed hormonal quantification and data analysis. All remaining experimental work was performed by Z.N.V. with help from C.M., W.L. and A.S. and supervision from J.H.. Z.N.V. and J.H. analysed the data, wrote the manuscript draft and incorporated feedback from all authors.

## Acknowledgements

We thank the Gatsby Charitable Foundation for funding Zoe Nemec Venza’s PhD. We thank Joe Cammarata, Adrienne Roeder and Mike Scanlon for the gift of the *Ppclv1a1brpk2* mutant and useful discussions about our work. We thank Lucia Primavesi for technical support and training. We thank Hana Martínková, Petra Amakorová and Kamila Wisnerová for their help with plant hormone analyses. The work was supported from ERDF project “Plants as a tool for sustainable global development” (No. CZ.02.1.01/0.0/0.0/16_019/0000827).

---

**Figure S1:**
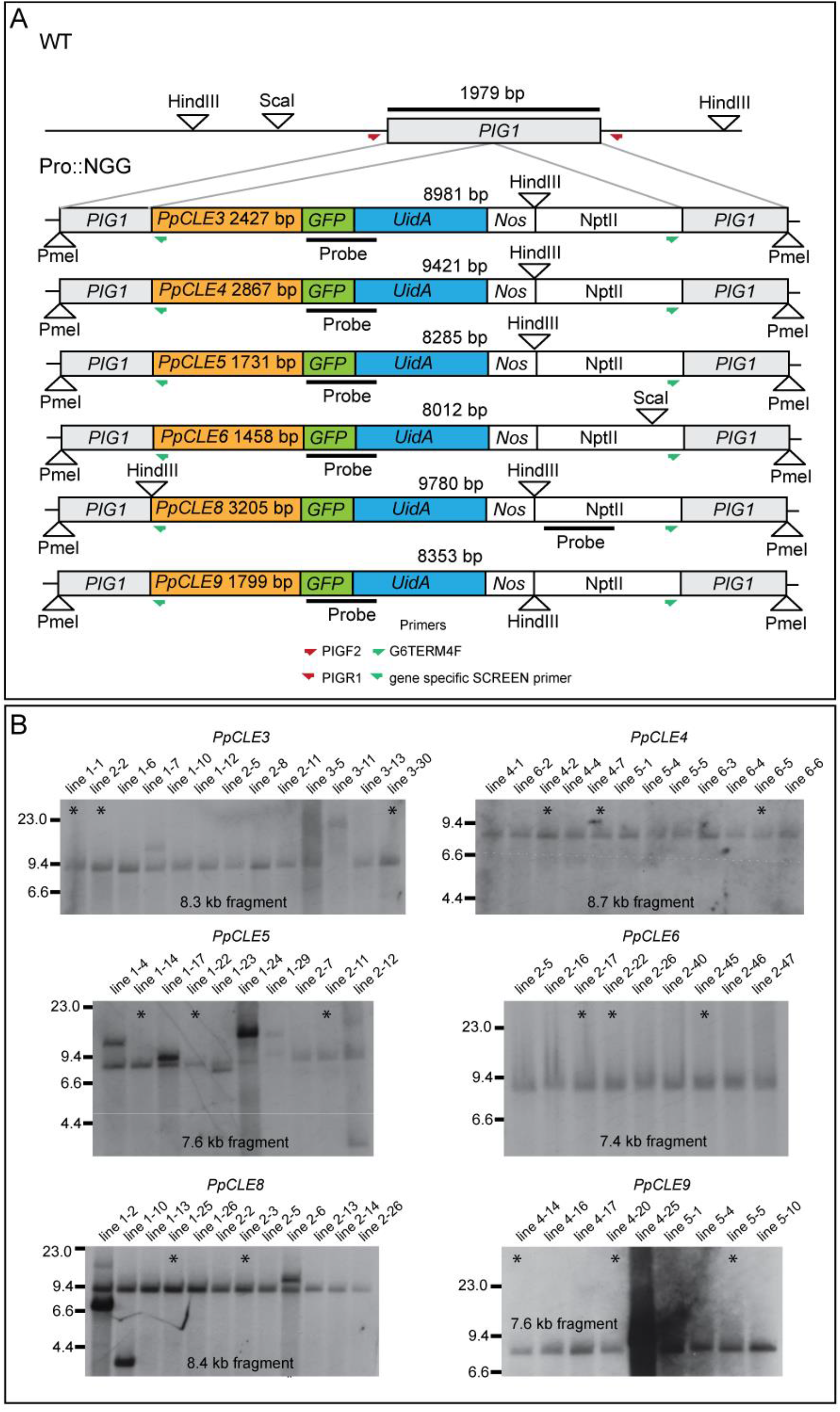
Strategy for generation of *promoter∷NLSGUSGFP* reporter lines. **(A)** Promoter fragments of varying lengths and including 5’UTRs of *PpCLE3*, *PpCLE4*, *PpCLE5* and *PpCLE6* were PCR amplified and cloned into the SmaI site of the PIG1NGGII [39] vector with an NptII casette in place of the BSD cassette. For *PpCLE8* and *PpCLE9* reporter lines, the first few amino acids of the coding sequence were also PCR amplified with promoter fragments. Orange boxes represent promoter fragments, *Nos* represents the *Nos* terminator and NptII represents the *CaMV35S-NptII-CaMVter* resistance cassette used to select positive transformants. Constructs were delivered into plants following linearization with PmeI, and lines were first screened by PCR using PIGF2 and promoter-specific primers or PIGR2 and G6TERMF primers [25] (data not shown). **(B)** PCR screening was followed by Southern analysis with a vector-specific probe following HindIII or ScaI digestion. Similar expression patterns were verified in three or more independently generated targeted insertants and lines labelled with asterisks were used in further expression analyses.

**Figure S2:**
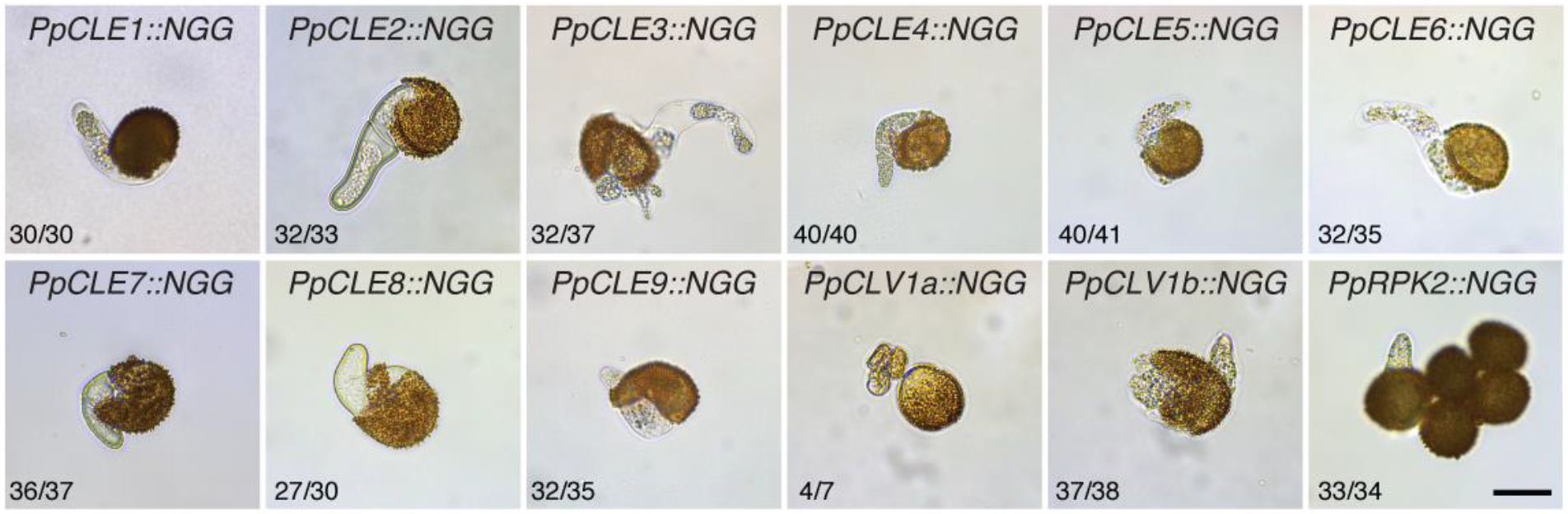
No CLAVATA expression was detected in germinating spores of *promoter∷NLSGUSGFP* reporter lines. Micrographs of germinating GUS-stained spores. The numbers in each panel indicate the proportion of sporelings displaying a similar expression pattern. Scale bar = 20 μm.

**Figure S3.**
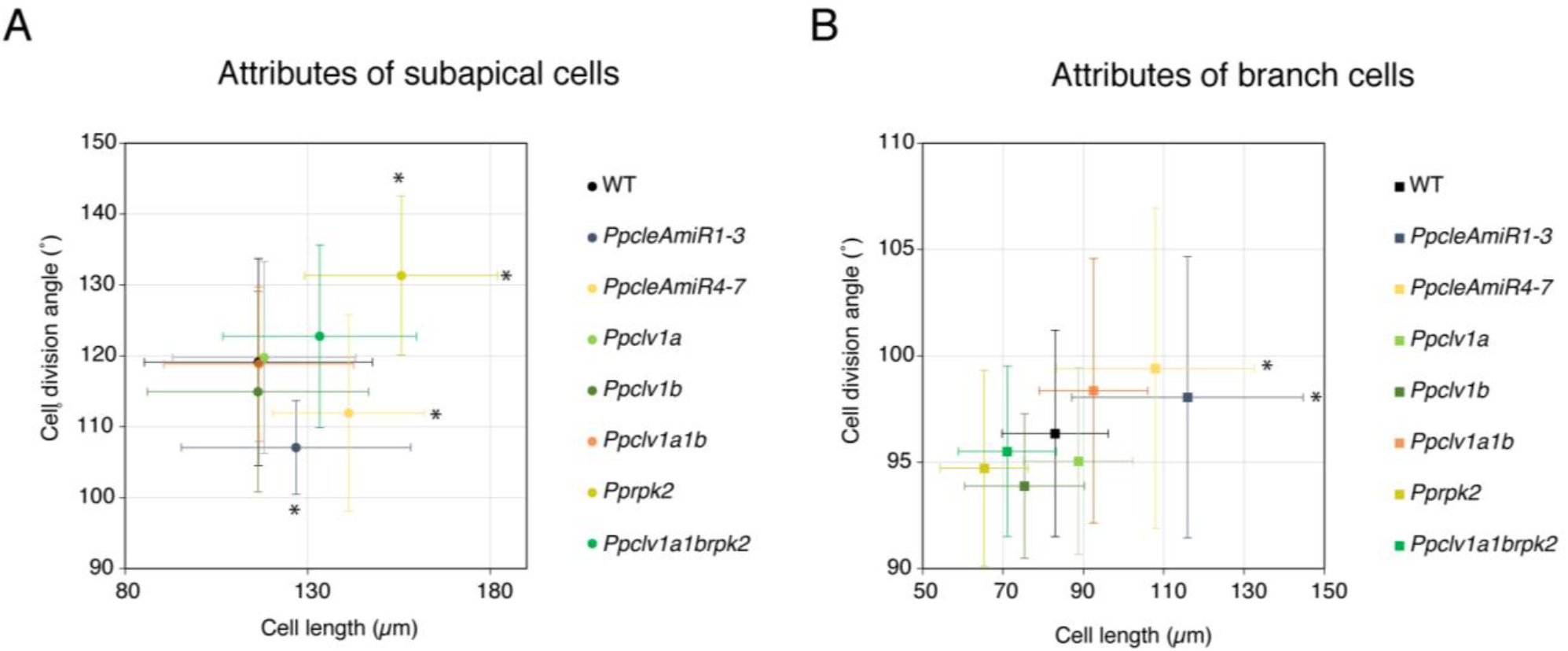
*PpcleAmiR1-3*, *PpcleAmiR4-7* and *Pprpk2* mutants have sub-apical cell length and division plane orientation defects in protonemata. **(A)** Analysis of caulonemal sub-apical cell attributes showed that *Pprpk2* cells are longer and with more oblique cell division planes than WT cells, *PpcleAmiR1-*3 mutants have longer cells and *PpcleAmiR4-7* mutants cells have divisions that are less oblique. *Ppclv1a1brpk2* subapical cells measurements are intermediate between wild-type and *Pprpk2* measurements. **(B)** Analysis of chloronemal sub-apical cell attributes showed that *PpcleAmiR1-*3 and *PpcleAmiR4-7* mutants have longer cells than WT. Bars indicate standard deviation. n ≥ 28, One-way ANOVA and Tukey test; * = p value < 0.05.

**Figure S4:**
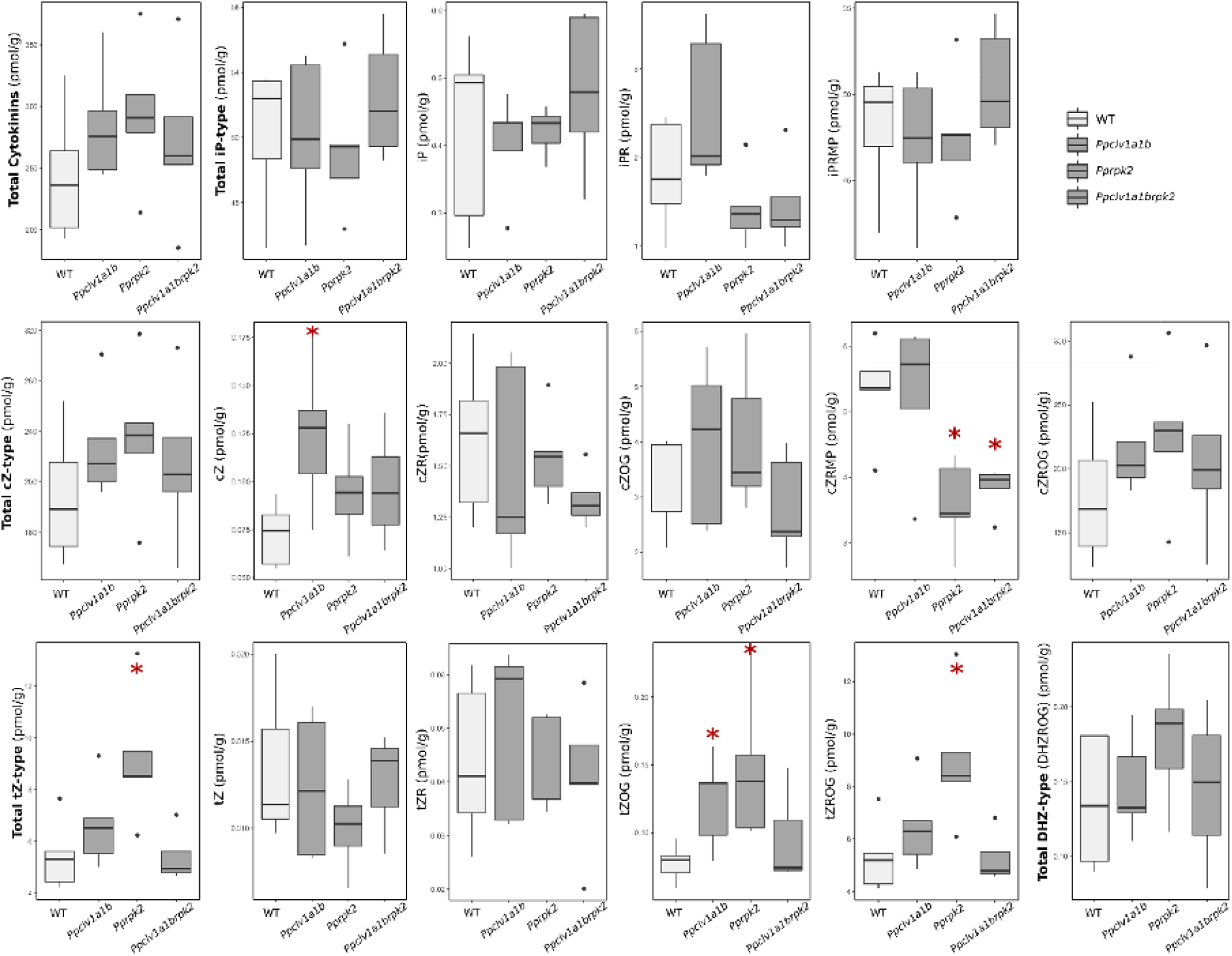
No difference in overall cytokinin levels between wild-type and *clavata* mutant lines was found. While total cytokinin and iP-type cytokinin content was similar between wild-type and mutant plants, cZ content was higher in *Ppclv1a1b* samples and cZRMP content was lower in *Pprpk2* and *Ppclv1abrpk2* samples. tZOG content was higher in *Ppclv1a1b* and *Pprpk2* mutants than in wild-type plants, with the latter also having higher tZROG. As a result, total tZ-type cytokinins content was higher in *Pprpk2* mutants than in wild-type plants. iP = N^6^-(Δ^2^-isopentenyl)adenine; iPR = N^6^-(Δ^2^-isopentenyl)adenosine; iPRMP = N^6^-(Δ^2^-isopentenyl)adenosine 5′-monophosphate; cZ = cis-zeatin; cZR = cis-zeatin riboside; cZOG = cis-zeatin O-glucoside; cZRMP = cis-zeatin riboside 5′-monophosphate; cZROG = cis-zeatin riboside O-glucoside; tZ = trans-zeatin; tZR = trans-zeatin riboside; tZOG = trans-zeatin O-glucoside; tZROG = trans-zeatin riboside O-glucoside; DHZROG = dihydrozeatin riboside O-glucoside. Other cytokinins were not detected. n = 5.

